# The effects of intense light on *Caenorhabditis elegans*

**DOI:** 10.1101/2025.06.05.658132

**Authors:** Marina Kniazeva, Gary Ruvkun

## Abstract

*Caenorhabditis elegans* utilize light receptors to modify foraging and locomotion in response to phototoxic blue and UV light. Our study investigated physiological and behavioral changes in *C. elegans* during and after exposure to high-intensity light-emitting diode (LED) light. Our findings corroborate previously reported blue-light-induced oxidative stress, mitochondrial damage, and avoidance behavior. Novel to our work is the identification of a protective role for lysosome-related organelles (LROs), the observation of a unique “shelter-seeking behavior” potentially mediated by light gradient sensing, and the seemingly dispensable role of canonical light receptors in this specific behavioral response.

## Introduction

*Caenorhabditis elegans*, a ubiquitous free-living nematode inhabiting diverse soil environments, naturally encounters a range of light conditions. The duration and intensity of this exposure fluctuate considerably due to circadian and seasonal cycles. Furthermore, environmental humidity introduces an additional layer of complexity, as water droplets on the nematode’s cuticle or surrounding substrate can either offer protection by scattering light or, conversely, potentially intensify its effects by acting as magnifying lenses, thereby influencing the actual light dose received. This dynamic interplay underscores the importance of understanding how *C. elegans* senses and responds to light. While it is known that *C. elegans* possesses light-sensing capabilities and exhibits behavioral responses to light ^1 2 3 4^, the full spectrum of physiological and behavioral adaptations to varying light conditions remains to be discovered.

Here we show that lysosome-related organelles, LROs, play a role in response to intense light exposure, a “shelter-seeking behavior” possibly driven by light gradient sensing and suggest that previously discovered light reception pathways are surprisingly not essential for this particular behavior.

## Results

### Exposure to intense light Increases *C. elegans* intestinal autofluorescence

The intestinal cells of *C. elegans* contain spherical granules known as lysosome-related organelles (LROs)^5^. LROs store and release various substances, including proteins and lipids, and function in zinc homeostasis, detoxification, and stress responses ^6, 7, 8^. While LROs exhibit green autofluorescence under UV excitation at high magnification, which can often interfere with GFP-tagged protein expression studies in the intestine, this autofluorescence is typically insignificant or invisible in healthy wild-type (N2 strain) larvae and young adults when viewed at low magnification.

We observed that a 1-hour exposure of wild-type adults, cultured on standard agar plates seeded with *E. coli* (OP50-2), to direct LED light at 137000 LUX intensity significantly enhances gut granule autofluorescence, making it detectable even under low magnification on a dissecting fluorescent microscope (Fig. 1A and 1B). Longer exposure overnight caused even more intense autofluorescence, highlighting the entire cytoplasm of intestinal cells, suggestive of granule disruption (Fig. 1C). Using blue and red color filters we found that the autofluorescence is triggered by the blue light component of the LED light (Fig. 1D and 1E).

**Fig. 1.**
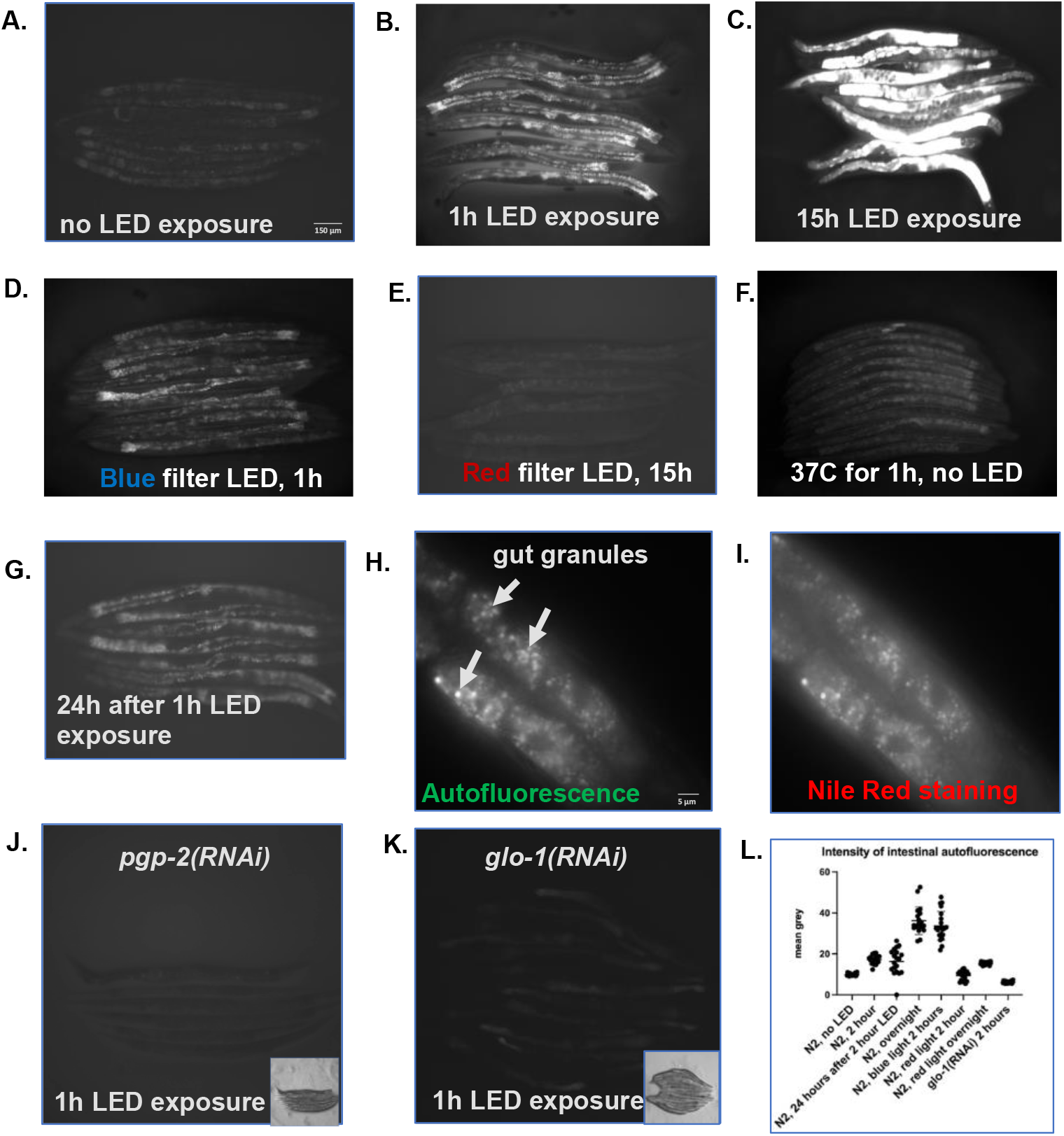
Exposure to LED promotes intestinal autofluorescence due to the blue light component. **A-F**, GFP images of wild type (N2) young adult animals exposed to LED or high temperature. **A**, unexposed control. **B** and **C**, exposure to LED for 1 and 15 h. **D** and **E**, exposure to blue (1h) and red (15h) emissions of LED. **F**, exposure to higher-than-normal temperature did not cause autofluorescence. **G**, Increased autofluorescence remains visible through the rest of lifespan. **H** and **I**, High magnification images of fluorescent gut granules ina wild type animal exposed to LED for 1 hour. **H**, autofluorescence is visible in GFP channel. **I.** Same granules are positive for the live dye, Nile Red, in RFP channel. **J** and **K**, *pgp-2(RNAi)* and *glo-1(RNAi)* RNAi treatment that prevents LRO formation inhibits LED induced intestinal autofluorescence. Scale bar 150 μm corresponds to images **A-G** and **J** and **K**, scale bar 5 μm corresponds to images **H** and **I.L**, Autofluorescence intestinal granules intensity calculated for A-G, mean grey.

While the overall temperature increase from an LED might be small, there could be localized heating at the surface of the agar where the LED light is focused. We addressed this concern by including a control group exposed to 37°C for 1 hour, a temperature significantly above the normal *C. elegans* culture conditions (Fig. 1F). These control animals did not show the same autofluorescence, suggesting that heat is not the trigger, and that the intense light (specifically the blue light component) causes the autofluorescence.

The LED induced autofluorescence persisted after removal from LED exposure (Fig. 1G), suggesting a lasting, or irreversible, change within the LROs.

Nile Red, a lipophilic dye, stains at least some LROs ^9^. We found that these Nile Red-positive granules are the same organelles exhibiting increased autofluorescence upon LED exposure (Fig. 1H and 1I). This intestinal autofluorescence in untreated animals is observable under high magnification in GFP channel. Prior genetic screens identified several genes essential for LRO formation^10, 11^. Using RNAi against two of these genes, *glo-1* and *pgp-2*, we demonstrated that such RNAi-treated animals did not develop autofluorescence during LED exposure (Fig. 1J and 1K). The intensity of intestinal autofluorescence after LED exposure was quantified (Fig. 1L). These findings confirm that the prominent LED-induced autofluorescence is localized to LROs.

A recent study reported rapid accumulation of autofluorescent material in LROs during exposure to the toxic compound benzaldehyde, suggesting that benzaldehyde is converted into autofluorescent products within LROs, contributing to detoxification^8^. In our case, the observed autofluorescence likely originates from LED-induced modifications within the LROs themselves.

### Exposure to LED Light Impairs Motility in *C. elegans*

Overnight exposure to LED light induces complete paralysis in wild-type *C. elegans*. This immobility is partially reversible; after a two-hour recovery period without LED treatment, the worms regain some movement, responding to touch and UV light with undulations. We investigated the potential causes of this paralysis, beginning with mitochondrial stress. Within two hours of LED exposure, we observed a significant increase in the expression of the mitochondrial stress reporter *cyp-14A4::pGFP* ^12^ (Fig. 2A), suggesting mitochondrial dysfunction. Using the *myo-3::GFP* reporter, we detected evidence of mitochondrial damage as early as one hour after the start of LED exposure (Fig. 2B). While these mitochondrial alterations likely contribute to the observed locomotion defects, they may not be the sole cause. Upregulation of *myo-3::GFP* has been previously reported in response to milder fluorescent light treatments that do not result in such severe locomotion impairment ^3^. This study also demonstrated that visible light induces photooxidation and subsequent upregulation of oxidative stress response enzymes. Therefore, it is reasonable to expect increased oxidative stress in our LED-exposed worms.

**Fig. 2.**
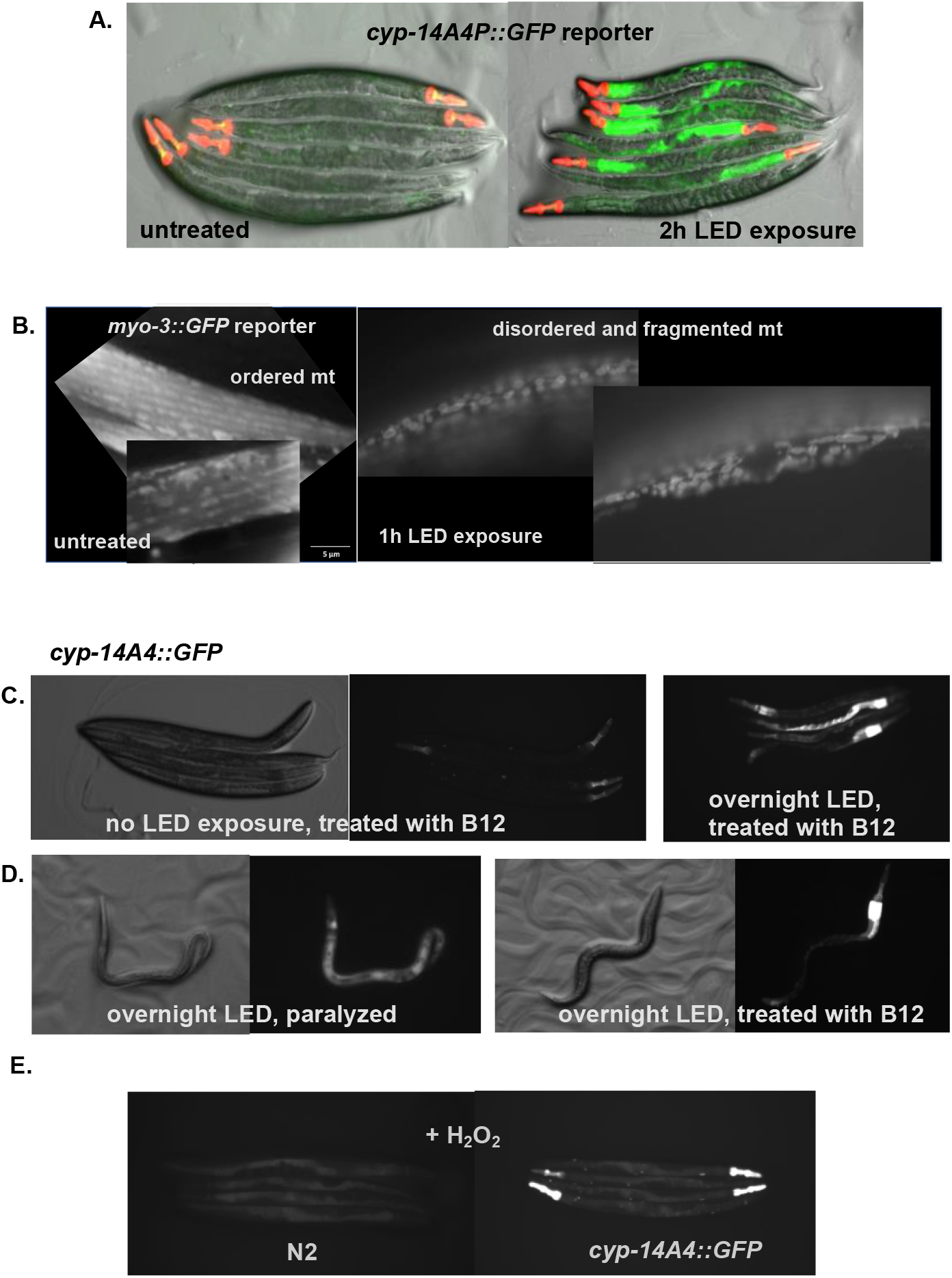
LED-induced activation of mitochondrial stress response. **A**, *cyp-14A::GFP* induction after 2h LED exposure. **B**, *myo-3::RFP* showing mitochondrial morphology in control (left panel) and LED-exposed worms (right panel). **C**, No *cyp-14A::GFP* expression in vitamin B12-treated worms under normal conditions. **D**, Partial inhibition of LED-induced *cyp-14A::GFP* by vitamin B12 (compare to Fig. 2A). **E** and **F**, DIC and GFP images of *cyp-14A::GFP* worms: **E**, paralyzed after overnight LED exposure and **F**, protected from paralysis by supplementation with vitamin B12. **G**, exposure to hydrogen peroxide does not induce *cyp-14A::GFP*.

To explore the role of oxidative stress, we investigated whether the antioxidant vitamin B12 ^13^ could mitigate the observed locomotion impairment. Supplementation with the B12 analog methylcobalamin (0.25 ng/ml, 600 µl per plate) significantly rescued locomotion in *cyp-14A4::pGFP* animals exposed to LED light overnight: 17 out of 20 B12 supplemented worms regained movement, compared to 0 out of 20 without supplementation (Fig. 2C and 2D). This suggests that oxidative stress plays a key role in the LED-induced paralysis. Treatment of young adult worms with hydrogen peroxide which generates reactive oxygen species (ROS) did not cause autofluorescence in wild type worms and did not up-regulate the *cyp-14A::pGFP* reporter (Fig. 2E).

While a shorter, two-hour LED exposure did not significantly affect locomotion, we did observe evidence of compromised mitochondrial integrity within this timeframe (Fig. 2B), indicating that mitochondrial damage precedes the overt paralysis.

We also investigated the role of lysosome-related organelles (LROs) in the intense light response. Intriguingly, while wild-type worms continued to forage on the bacterial lawn during short high intensity LED exposure, *pgp-2* and *glo-1* mutants, which lack LROs, exhibited pronounced avoidance behavior (Fig. 3A and 3B). Moreover, unlike wild-type worms, most *pgp-2* and *glo-1* mutants exposed to LED light for just two hours were found dead the following day (Fig. 3C). These findings strongly suggest that LRO-deficient mutants are more susceptible to the toxic effects of LED exposure, implicating a protective role for LROs against intense light-induced damage.

**Fig. 3.**
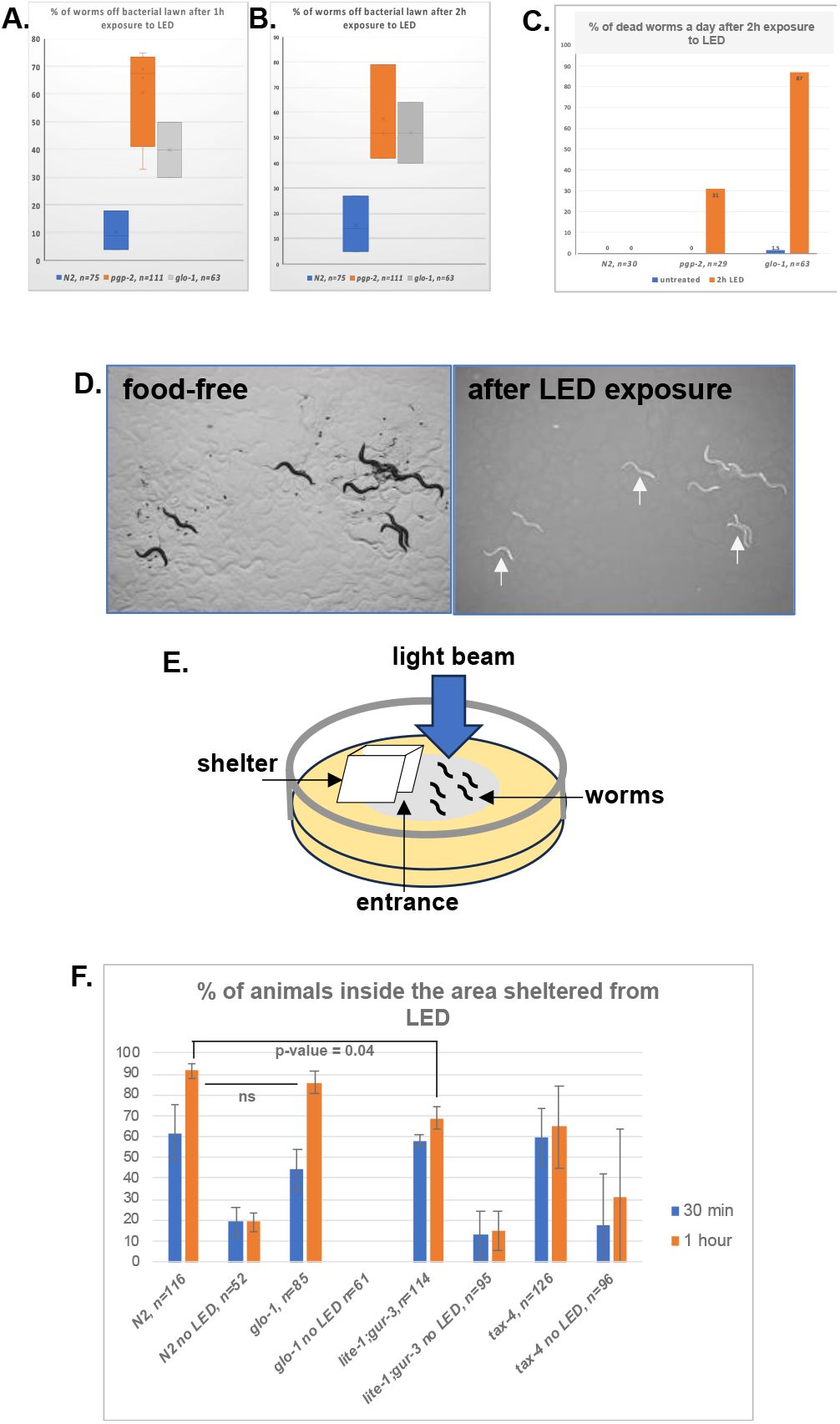
Animals lacking LROs are more sensitive to the LED exposure. **A** and **B**, Avoidance assay: percentage of wild type, *glo-1*, and *pgp-2* animals outside the bacterial lawn after 1 (A) and 2 hours (B) of LED exposure. **C**, Survival assay: percentage of dead animals of indicated genotypes 24 hours post 2-hour LED exposure. **D**, Autofluorescence in recovering worms: DIC and GFP channel images of wild-type animals after LED-induced burrowing on food-free plates, with arrows pointing to autofluorescent bodies. **E**, Experimental setup for shelter-seeking assay: Cartoon depicting the aluminum foil shelter, agar, bacterial lawn, and LED light. **F**, Shelter-seeking assay: Percentage of animals of each genotype within the shelter after 30 minutes and 1 hour of LED exposure.

### Exposure to intense light activates shelter seeking behavior in *C. elegans*

To distinguish between food avoidance and the escape behavior, we tested if the worms exposed to intense LED light would seek shelter or disperse randomly. We found that in the absence of food (regular plates w/o bacterial spot), 90% (n=20) of wild type worms burrowed inside the agar within 1h after LED exposure and all the worms re-appeared on the surface of agar within 2 hours after the light went off. No burrowing behavior was observed in the presence of food or in control animals on food-free plates without LED exposure during the same period. These results suggested that leaving the lawn is likely to be an escape behavior rather than food avoidance. Surprisingly, the following day all the worms on a food-free plate were found dead or sick and immobilized, while control untreated with intense light animals on food-free plates were alive and mobile. This suggested that either feeding protects from intense light induced toxicity or 1-hour intense light exposure prevents survival without food (Table 1). The worms exposed to intense light for 1 hour on food-free plates do have increased autofluorescence (Fig. 3D). This observation further supports the previous suggestion that this autofluorescence originates internally and is not derived from food.

**Table 1.** % of worms inside agar on food-free plates (burrowing behavior), n=40.

| **LED exposure, 1h** | **post LED, 2h** |
| --- | --- |
| **90%** | **0%** |

We also tested if the animas exposed for intense light would actively seek an escape route. We’ve set up a small shelter (cubicle ∼ 1×1 cm) made of aluminum foil covering part of the bacterial lawn with one entrance facing the lawn (Fig. 3E). We found that wild type worms crowded inside the sheltered area within 30 min after intense light exposure and remained there until the light went off (after 1 hour of expose) (Fig. 3F). A number of animals under the roof was significantly low on the control plates with the shelter not exposed to the light (p-value = 1.5854E-05). Suggesting that a presence of the aluminum structure did not attract the worms in the absence of intense light. We also found that worms lacking LRO, *glo-1*, behave similarly to wild type sheltering during LED exposure and ignore the covered area randomly dispersing on the plate (Fig. 3E). This suggests that short exposure to LED light induces active shade seeking behavior and is unrelated to changes in LRO increased autofluorescence.

### Light receptors are not necessary for LED activated behavior

There are two known light receptors in *C. elegans* encoded by *lite-1* and *gur-3* genes ^1^. Double mutants have almost no response to blue/UV light ^1^. We used the double mutant, *lite-1;gru-3*, to test if these receptors mediated the shade seeking behavior similarly to photophobic response to UV light ^14^.

We did not detect a difference between behavior of wild type and *lite-1;gru-3* double mutant animals at 30 min and only a small difference (p-value=0.04) at 1 hour (Fig. 3F). These results indicated that neither of the receptors are necessary for initiation and maintenance of LED light shelter seeking behavior.

To rule out temperature gradients as a potential factor in shelter-seeking behavior during LED exposure, we tested *tax-4* null mutants ^15^, which are unable to respond to temperature gradients. We found that these mutants accumulated inside the shelter only slightly less than wild-type animals (p-value = 0.01) (Fig. 3F). This result suggests that temperature variation if any on the plate did not significantly contribute to the observed behavior.

## Discussion

The impact of visible light exposure on *C. elegans* has been extensively studied, notably by the Dillin group ^3^. Their work demonstrated that prolonged exposure to visible (blue) light leads to a shortened lifespan, increased production of reactive oxygen species (ROS), activation of the unfolded protein response (UPR), and mitochondrial damage. Initially unaware of these findings, our project began with the independent goal of investigating whether *C. elegans* could emit light. This led us to use a high-intensity LED light source, suitable for our intended purpose. Upon analyzing our initial data, and as newcomers to the field, we began to compare our observations with existing published results.

Consistent with the findings of Dillin and colleagues ^3^, our data obtained with a different set of reagents also revealed evidence of oxidative toxicity and mitochondrial damage following high-intensity LED exposure. Furthermore, our investigation uncovered a protective role for lysosome-related organelles (LROs). Notably, *C. elegans* LROs have been recently implicated in cytoprotective stress responses, detoxification processes, the activation of immune effector genes ^8^. Aligning with these established functions, we observed that LROs play a protective role during high-intensity LED light exposure. This raises questions about the underlying mechanisms of this specific protection. Future research could explore potential mechanisms such as detoxification or recycling of damaged molecules, membrane repair, and/or other processes. Additionally, identifying the specific LRO components involved in this protection would be valuable avenues for future investigation.

We also uncovered a peculiar “shelter-seeking behavior” that appears distinct from simple avoidance responses, where animals typically disperse randomly away from aversive stimuli, such as feeding on toxic bacteria^16^ or burrowing into agar in the absence of food as previously described. Notably, we observed no burrowing behavior in the presence of bacterial food, indicating that the attraction to the food source overrides the potential danger of phototoxicity. Intriguingly, even when food was available, the worms actively moved towards a shaded area, which we termed a “shelter,” suggesting a possible ability to sense and respond to a light gradient, analogous to chemo- and thermotaxis. Surprisingly, the two known light receptors, *lite-1* and *gru-3*, responsible for mediating many light-induced phenotypes in *C. elegans* were found to be dispensable for this specific “shelter-seeking” behavior. This raises a question: if canonical light receptors are not involved, what alternative sensory mechanisms might be at play? One possibility is that other tissues in *C. elegans* possess a more general photoreceptive capacity that mediates this response.

## Methods

### Worm strains

wild type N2 Bristol, *glo-1(zu391), pgp-2(kx48), lite-1(ce314), lite-1(ce314);;gur3-(ok2245), tax-4(p678), myo-3::GFP syIs243 [myo-3p::TOM20::mRFP + unc-119(+) + pBS Sk+]* were obtained from the Caenorhabditis Genetic Center (CGC). *cyp-14A4P::GFP* was generated in the laboratory by Kai Mao ^12^.

### Worm plates

Standard 6 cm Nematode Growth Media (NGM)^17^ agar plates were seeded with 300 µl *E. coli* OP50-1 cultured overnight at 37ºC.

### RNAi treatment

*C. elegans* adults were bleached and eggs were plated on bacterial lawn seeded with HT115 E. coli expressing RNAi from Ahringer’s RNAi library. Each RNAi agar plate was 6 cm in diameter, contained 50 μg/ml ampicillin, and 5 mM IPTG and was seeded with 300 μl of individual overnight HT115 RNAi cultures grown in liquid LB supplemented with 50 μg/ml ampicillin^18^.

### LED exposure

LED source Microscope LED Illuminator MP-LED150 equipped with Volpi 1900 fiber optic light guide cable. The fiber optic output was positioned 3 cm above the agar surface at room temperature illuminating to approximately 137000 LUX. Illumination was measured with LED Light Meter LT40 (EXTECH Instruments). Color filters were obtained from Arbor Scientific (33-0190). Red filter illumination: 6750 LUX, blue filter illumination: 2672 LUX. Young adult worms (typically 30-50) were exposed to the LED light for indicated periods of time.

### Heat shock

For heat shock NGM plates with young adult worms were incubated at 37ºC for 1 hour.

### Hydrogen peroxide treatment

50 µL of 3% hydrogen peroxide (Sigma-Aldrich, 88597) was added to bacterial lawn with young adult animals so that all of the lawn area was covered. An appearance of bubbles (oxygen) indicated the presence of hydrogen peroxide and catalase activity: 2H_2_O_2_ →catalase→ 2H_2_O+O_2_. The plates were kept at room temperature. The treated animals were evaluated on the plates under UV microscope immediately and with 30 min increments for 6 hours.

### Nile red staining

young adults exposed to LED were transferred to a new NGM plate in a drop of NaN3 (100 mM) in the presence of Nile Red (Invitrogen, N1142) at 10 μg/ml in M9 stain.

Vitamin B12 treatment: The vitamin B12 stable analog, Methylcobalamin (Sigma M9756), was added directly to the plates at 0.25 ng/ml, 600 µl per plate.

### Avoidance assay

The experimental set up was done as described. 30-60 worms were placed on bacterial lawn under the LED beam. Number of worms inside the shelter were counted in 30 min or 1 hour after the exposure. The assay was done in two biological replicates.

Quantitative analyses were performed in Microsoft Excel and Prism 10 for macOS.

### Microscopy

Live *C. elegans* were visualized with Nicon SMZ18, AxioZoom V16 (Zeiss) and AxioImager Z.1 (Zeiss) microscopes. Zeiss microscopes were equipped with OrkaFlash 4.0 (Hamamatsu) and Axiocam HRc (Zeiss) cameras, respectively. Images were processed with ImageJ software.

## Acknowledgments

We thank WormBase for genome information and curation. Some strains were provided by the Caenorhabditis Genetics Center, which is funded by NIH Office of Research Infrastructure Programs (P40 OD010440). This work was supported by the NIH National Institute on Aging R01AG16636 and R01AG043184 to G.R.

## Notes

### Competing Interest Statement

The authors have declared no competing interest.

